# Characterisation and comparison of semen microbiota and bacterial load in men with infertility, recurrent miscarriage, or proven fertility

**DOI:** 10.1101/2024.02.18.580923

**Authors:** Shahriar Mowla, Linda Farahani, Tharu Tharakan, Rhianna Davies, Gonçalo D S Correia, Yun S Lee, Samit Kundu, Shirin Khanjani, Emad Sindi, Raj Rai, Lesley Regan, Dalia Khalifa, Ralf Henkel, Suks Minhas, Waljit S Dhillo, Jara Ben Nagi, Phillip R Bennett, David A MacIntyre, Channa N Jayasena

## Abstract

Several studies have associated seminal microbiota abnormalities with male infertility but have yielded differing results owing to their limited sizes or depths of analyses. The semen microbiota during recurrent pregnancy loss (RPL) has not been investigated. Comprehensively assessing the seminal microbiota in men with reproductive disorders could elucidate its potential role in clinical management. We used semen analysis, terminal-deoxynucleotidyl-transferase-mediated-deoxyuridine-triphosphate-nick-end-labelling, Comet DNA fragmentation, luminol ROS chemiluminescence and metataxonomic profiling of semen microbiota by16S rRNA amplicon sequencing in this prospective, cross-section study to investigate composition and bacterial load of seminal bacterial genera and species, semen parameters, reactive oxidative species (ROS), and sperm DNA fragmentation in men with reproductive disorders and proven fathers. 223 men were enrolled included healthy men with proven paternity (n=63), the male partners in a couple encountering RPL (n=46), men with male factor infertility (n=58), and the male partners of couples unexplained infertility (n=56). Rates of high sperm DNA fragmentation, elevated ROS and oligospermia were more prevalent in the study group compared with control. In all groups, semen microbiota clustered into three major *genera*-dominant groups (1, Streptococcus; 2, Prevotella; 3, Lactobacillus and Gardnerella); no species clusters were identified. Group 2 had the highest microbial richness (P<0.001), alpha-diversity (P<0.001), and bacterial load (P<0.0001). Overall bacterial composition or load has not found to associate with semen analysis, ROS or DNA fragmentation. Whilst, global perturbation of the seminal microbiota is not associated with male reproductive disorders, men with unidentified seminal *Flavobacterium* are more likely to have abnormal seminal analysis. Future studies may elucidate if *Flavobacterium* reduction has therapeutic potential.

## Introduction

Mean sperm counts reported within clinical studies have reduced annually by 2.6% since 2000 (1). Male factor accounts for approximately half of all cases of infertility yet there are limited available interventions to improve sperm quality. Understanding the pathogenesis of male infertility may reveal novel therapeutic approaches for treating affected couples.

Symptomatic, genitourinary infection is an established cause of male infertility which may be detected by semen culture and treated with antibiotics (2, 3). The current European Association of Urology guidance states that whilst antibiotics may improve overall semen quality, there is no evidence of increased pregnancy rates after antibiotic treatment of the male partner (4) (5). Seminal leukocytes release bactericidal reactive oxygen species (ROS) in response to infection, however paradoxically this may damage sperm DNA and impair semen quality (6). We and others have reported that asymptomatic men affected by recurrent pregnancy loss (RPL), infertility or impaired preimplantation embryo development have increased risks of high seminal ROS and sperm DNA fragmentation (7, 8, 9, 10, 11, 12). It is therefore plausible that asymptomatic seminal infections may be associated with impaired reproductive function in some men. Since semen culture has a limited scope for studying the seminal microbiota due to its inability to identify all present microbiota next generation sequencing (NGS) approaches have been reported recently by a growing number of investigators (13, 14, 15, 16, 17, 18, 19). These studies, with varying methodologies, have produced inconsistent and conflicting results. As such, the composition of semen microbiota and its associations with clinical and molecular markers of male reproduction remains understudied. Elucidation of an association would have wide clinical application with therapeutic potential couple with reproductive disorders (20).

We hypothesised that semen microbiota composition associates with functional semen parameters, including ROS levels and sperm DNA fragmentation. To test this, we explored relationships between metataxonomic profiles of bacteria, bacterial copy number and key parameters of sperm function and quality in semen samples collected from 223 men, including those diagnosed with male factor infertility, unexplained infertility, partners affected by recurrent miscarriage, and paternity-proven controls.

## Methods

*Ethical approval* was granted by the West London and Gene Therapy Advisory Committee (GTAC) Research Ethics Committee (14/LO/1038) and by the Internal Review Board at the Centre for Reproductive and Genetic Health (CRGH) (IRB-0003C07.10.19). Participants were recruited following informed consent from clinics in Imperial College London NHS Trust and The Centre for Reproductive and Genetic Health (CRGH). Further detailed information on methods used in this study are included in the Supplementary Material.

*Semen samples* were produced by means of masturbation after 3-7 days abstinence. All semen samples were collected into sterile containers after cleaning of the penis using a sterile wipe. Samples were incubated at 37°C for a minimum of 20 mins prior to analysis. An aliquot was collected in a sterile cryovial and stored at −80°C.

*Diagnostic semen analysis* was carried out according to WHO 2010 guidelines and UK NEQAS accreditation (21) (22). Seminal analysis was performed in the Andrology Departments of Hammersmith Hospital and CRGH. Microscopic and macroscopic semen qualities were assessed within 60 mins of sample production. Semen volume, sperm concentration, total sperm count, progressive motility and total motility count, morphological assessment, anti-sperm antibodies and leucocyte count were established.

*ROS analysis* was performed using an in-house developed chemiluminescence assay validated by Vessey et al (23). Results are therefore reported as ‘relative light units per second per million sperm’. The upper limit of optimal ROS was internally determined at 3.77 RLU/sec/106 sperm (95% CI) (24).

*Sperm DNA fragmentation assessment* performed by TUNEL (Terminal deoxynucleotidyl transferase biotin-dUTP Nick End Labelling) assay defined elevated sperm DNA fragmentation as >20% (25). Samples for the COMET assay were sent to the Examen Lab (Belfast, UK) for analysis with elevated sperm DNA fragmentation defined as >27% (26).

*DNA extraction* was performed on 200 μL of semen using enzymatic lysis and mechanical disruption. Bacterial load was estimated by determining the total number of 16S rRNA gene copies per sample using the BactQuant assay (27).

*Metataxonomic profiling of semen microbiota* was performed using MiSeq sequencing of bacterial V1-V2 hypervariable regions of 16S rRNA gene amplicons 16S rRNA genes using a mixed forward primerset 28F-YM GAGTTTGATYMTGGCTCAG, 28F-Borrelia GAGTTTGATCCTGGCTTAG, 28F-Chloroflex GAATTTGATCTTGGTTCAG and 28F-Bifdo GGGTTCGATTCTGGCTCAG at a ratio of 4:1:1:1 with 388R reverse primers. Sequencing was performed on the Illumina MiSeq platform (Illumina, Inc. San Diego, California). Following primer trimming and assessment of read quality, amplicon sequence variants (ASV) counts per sample were calculated and denoised using the Qiime2 pipeline (28) and the DADA2 algorithm (29). ASVs were taxonomically classified to species level using a naive Bayes classifier trained on all sequences from the V1-V2 region of the bacterial 16S rRNA gene present in the SILVA reference database (release 138.1) (30) (31).

Further methodological detail can be found in supplementary text.

**Controls and contamination** 3 negative kit/environmental control swabs were included to identify and eliminate potential sources of contamination and false positives in the 16S m*etataxonomic profiles.* These swabs were removed from the manufacturers packaging, waved in air, and then subjected to the same entire DNA extraction protocol. Decontamination of data was done using the decontam package (v1.9.0) in R, at ASV level, using both “frequency” and “prevalence” contaminant identification methods with *threshold* set to 0.1 (31). The “frequency” filter was applied using the total 16S rRNA gene copies measured as the *conc* parameter. For the “prevalence” filter all 3 blank swabs were used as negative controls and compared against all semen samples. ASVs classified as a contaminant by either method (n = 94) were excluded.

### Statistical analysis

Hierarchical clustering with Ward-linkage and Jensen-Shannon distance was used to assign samples to putative community state types, with the number of clusters chosen to maximise the mean silhouette score. Linear regression models used to regress microbiota features against semen quality parameters and other clinical and demographic variables were fitted with the base R *lm* function (v4.2.0). The Benjamini-Hochberg false discovery rate (FDR) correction was used to control the FDR of each covariate signature independently (e.g., ROS, DNA Fragmentation, or Semen quality), with a q < 0.05, or 5%, cut-off, in both regression and Chi-squared analyses. Detailed information for statistical modelling is presented Supplementary methods.

## Results

### Study population

Semen samples were collected from a total of 223 men; this included control (n=63) and a study group (n=160) comprised of men diagnosed with male factor infertility (MFI) (n=58), male partners of women with recurrent pregnancy loss (RPL) (n=46) and male partners of couples diagnosed with unexplained infertility (UI) (n=26). The overall mean age of the total cohort was 38.1 ± 6 (mean ± SD). The mean age for controls was 40.1 ± 8, and the mean age for patients undergoing various fertility investigations was 37 ± 4.8. Ethnicity representation amongst recruited cohorts were not significantly different (p=0.38, Chi-square; Supplementary Table 1).

### Semen quality assessment

Rates of high sperm DNA fragmentation, elevated ROS and oligospermia were more prevalent in the study group compared with control (Table 1). The study group represented 85% of samples with high sperm DNA fragmentation, 85% of samples with elevated ROS and 79% of samples with oligospermia. Rates of abnormal seminal parameters including low sperm concentration, reduced progressive motility and ROS concentrations were found to be highest in the MFI group (Supplementary Figure 1). Baseline characteristics between the RPL, unexplained subfertility and controls groups were similar (Supplementary Figure 1)

**Table I:**
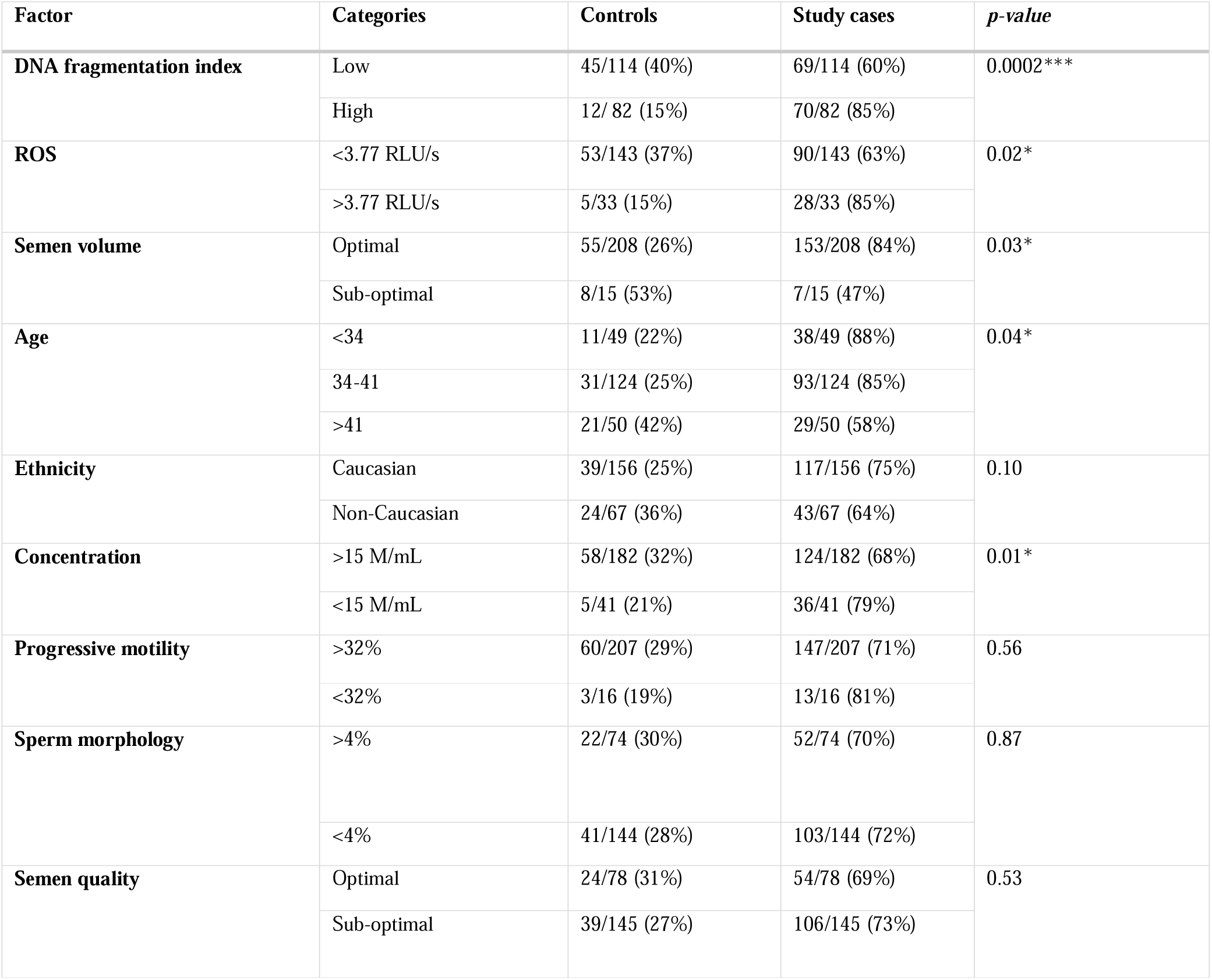
Patient demographics and notable parameters of seminal quality and function for controls and study subjects. Fisher’s exact tests for all except age. Chi-square test for age. (n=223).

### Seminal microbiota

Following decontamination, a total of 7,998,565 high quality sequencing reads were identified and analysed. Hierarchical clustering (Ward linkage) of relative abundance data resolved to genera level identified three major clusters, as determined by average silhouette score, amongst all samples (Figure 1, Supplementary Figure 2). These were compositionally characterised by highest mean relative abundances of 1. *Streptococcus* (23.8%), 2. *Prevotella* (24.4%), or 3. *Lactobacillus* and *Gardnerella* (35.9% and 35.6%, respectively, Figure 1C). Assessment of bacterial load using qPCR showed Clusters 2 and 3 had significantly higher bacterial loads compared to Cluster 1. Similar analyses were performed using sequencing data mapped to species level, however, examination of individual sample Silhouette scores within resulting clusters highlighted poor fitting indicating a lack of robust species-specific clusters (Supplementary Figure 3). To further investigate potential pairwise ecological interactions between taxa, a co-occurrence analysis was performed on the sequencing data, mapped to species level, with the SparCC algorithm (Figure 2). 5 major graph communities were detected. Community 1 highlighted a co-occurrence pattern between *Gardnerella vaginalis* and *Lactobacillus iners,* in agreement with the composition of cluster 3 from the hierarchical clustering analysis at genera level. Taxa belonging to communities 3 and 4 had a high number of connections (higher node degree), both within and between the two communities, including some anti-co-occurrence patterns (SparCC ρ < 0). These communities included species from Genera *Staphylococcus, Peptoniphilus, Corynebacterium, Prevotella,* among others.

**Figure 1.**
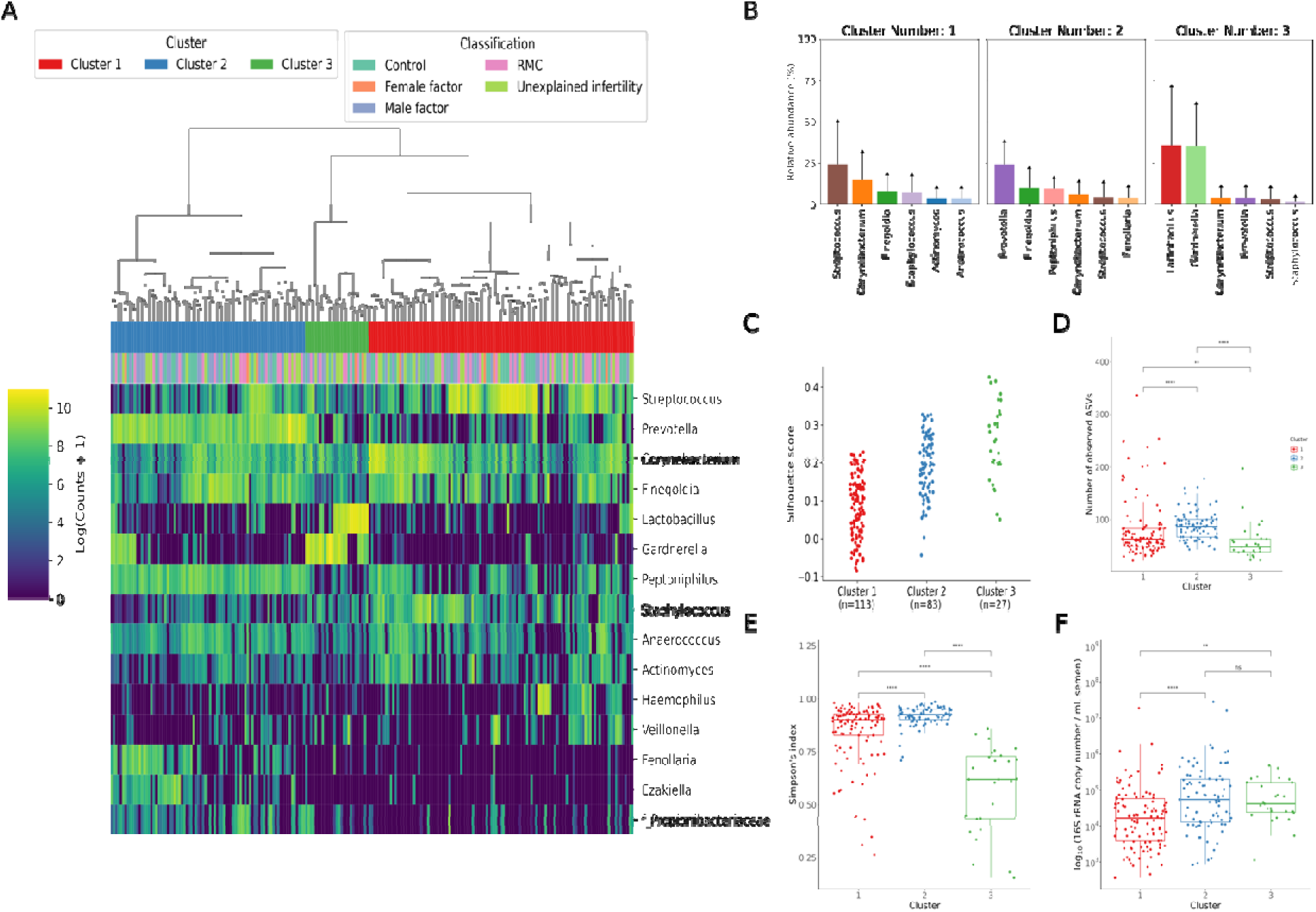
Characterisation of semen microbiota composition at genera level. **A)** Heatmap of Log10 transformed read counts of top 10 most abundant genera identified in semen samples. Samples clustered into three major microbiota groups based mainly on dominance by *Streptococcus* (Cluster 1), *Prevotella* (Cluster 2), or *Lactobacillus* and *Gardnerella* (Cluster 3). (n=223, Ward’s linkage). **B)** Relative abundance of the top 6 most abundant genera within each cluster. **C)** Silhouette scores of individual samples within each cluster. **D)** Species richness (p<0.0001; Kruskal-Wallis test) and **E**) alpha diversity (p<0.0001; Kruskal-Wallis test) significantly differed across clusters. **F**) Assessment of bacterial load using qPCR showed Clusters 2 and 3 have significantly higher bacterial loads compared to Cluster 1 Dunn’s multiple comparison test was used as a post-hoc test for between group comparisons (*p<0.05, ****p<0.0001).

**Figure 2.**
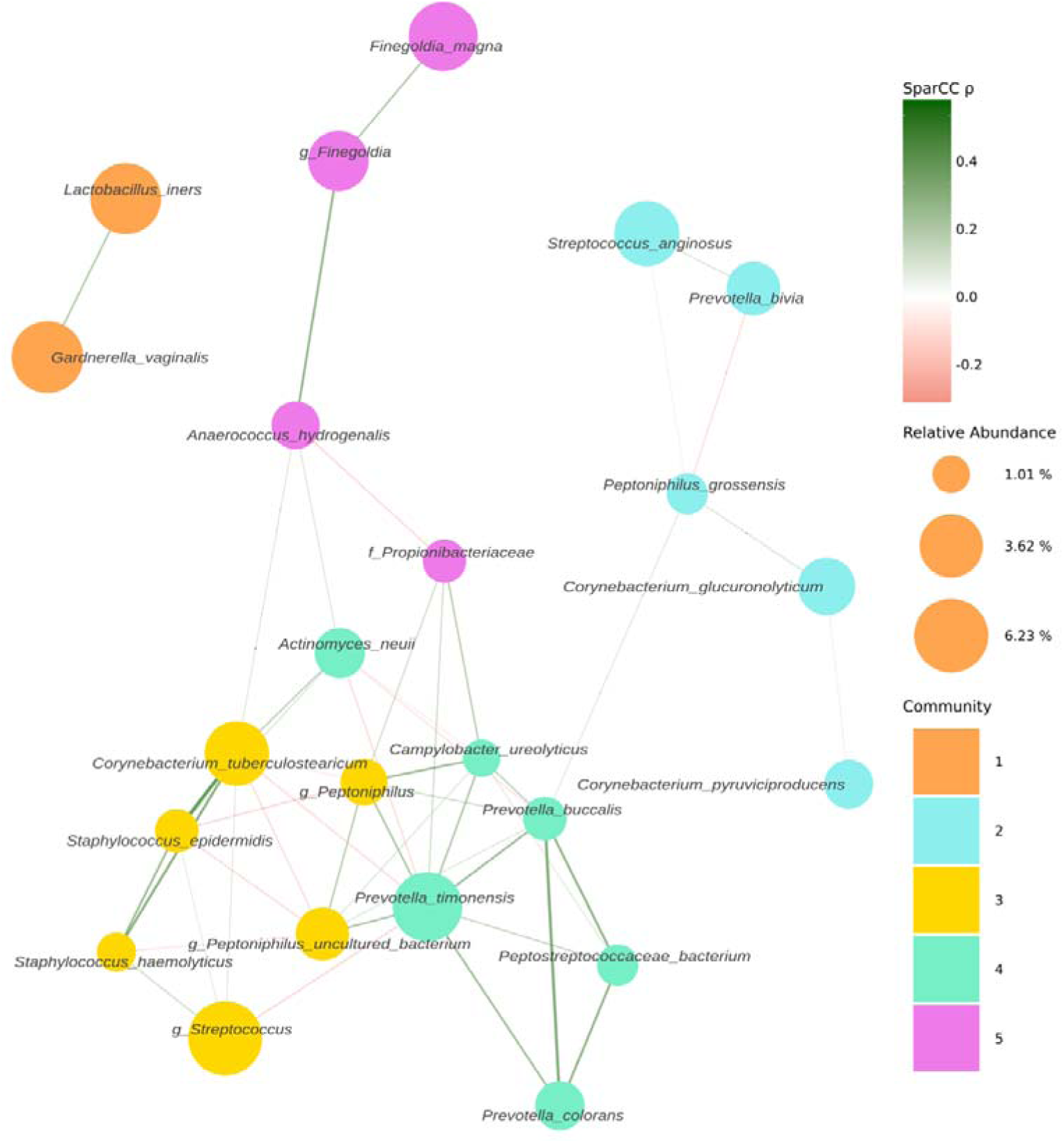
Co-occurrence network estimated with SparCC from 16S sequencing counts at species level. Network representing co-occurrence patterns (edges), between various taxonomic units, assigned at species level (nodes). Edges are colored by their estimated SparCC correlation coefficient (ρ). Edges with a SparCC bootstrapped *p-value* < 0.05, ρ < 0.25, and singleton nodes are not shown. Node color represents network community membership. Node sizes are proportional to the mean relative abundance of their respective taxon.

Bacterial richness, diversity and load were similar between all patient groups examined in the study (Supplementary Figure 4). Similarly, no significant associations between bacterial clusters, richness, diversity or load with seminal parameters, sperm DNA fragmentation or semen ROS were observed (Supplementary Tables 2-3). No significant differences in relative abundance of bacterial taxa between patient groups were detected, at genus or species-level. Several organisms at the genus level, identified variably in the literature as responsible for genito-urinary infection, were observed in our dataset but their prevalence did not reach our criteria (present in at last 25% of the samples) to be carried forward to regression modelling (32) (33) (24). This included *Chlamydia, Ureaplasma, Neisseria, Mycoplasma* and *Escherichia*. However, several associations (p<0.05) between relative abundance of specific bacterial genera and key sperm parameters were observed (Table 2). These included increased sperm DNA fragmentation was positively associated with increased relative abundance of *Porphyromonas* and *Varibaculum* and inversely correlated with *Cutibacterium* and *Finegoldia.* ROS was positively associated with *Lactobacillus* species relative abundance, with analyses performed at species level taxonomy indicating that this relationship was largely driven by *L. iners* (p=0.04; Table 3). In contrast, *Corynebacterium* was inversely associated with ROS and positively associated with semen volume. Of note, *Flavobacterium* genus was positively associated with both abnormal semen quality and sperm morphology and in both cases, withstood FDR correction for multiple testing (q=0.02 and q=0.01, respectively) (Table 2) (Figure 3). Consistent with this, a positive association between an unidentified species of *Flavobacterium* and semen quality was also observed (q=0.01, Table 3).

**Figure 3.**
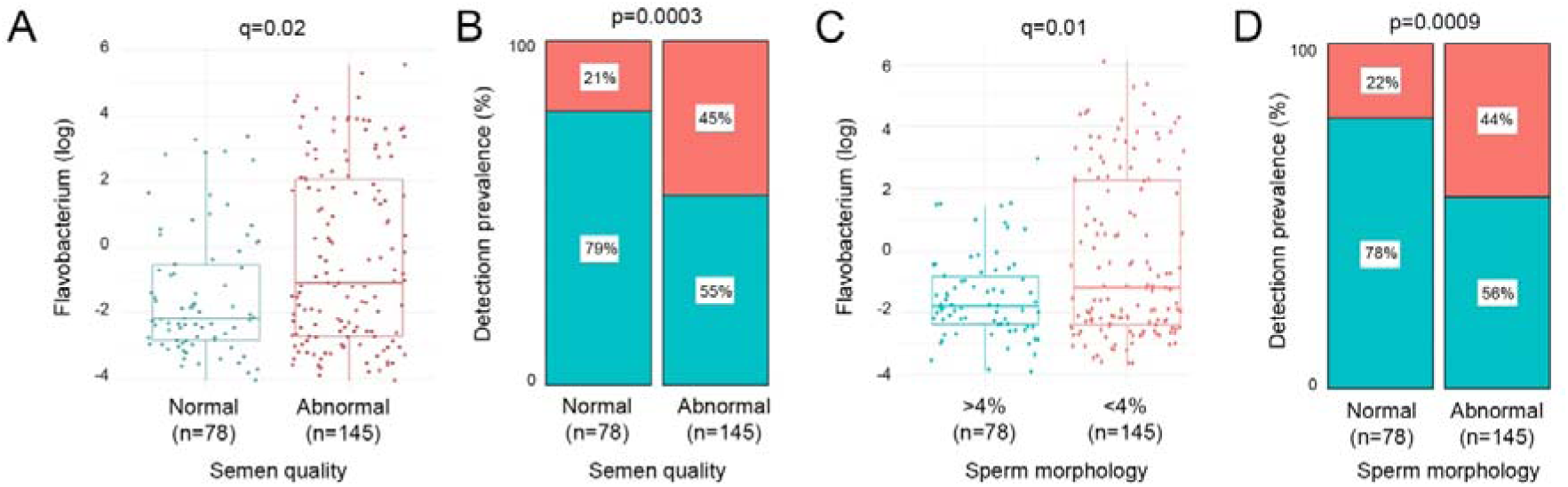
Relative abundance and prevalence matrices of Flavobacterium in relation to semen quality and morphology. **A)** Relative abundance of Flavobacterium was significantly higher in samples with abnormal semen (p=0.0002, q=0.02). **B)** Detection of flavobacterium was significantly more prevalent in abnormal semen quality samples (p=0.0003). **C)** Flavobacterium relative abundance was significantly higher in samples with <4% morphologically normal forms (p=0.0002, q=0.01). **D)** Flavobacterium was also significantly more prevalent in samples with low percentage of morphologically normal sperm (p=0.0009).

**Table II:** Differential abundance analysis for bacterial genera with seminal quality and functional parameters. Positive t-values indicate a positive relationship, and a negative t-value describes a negative relationship between relative abundance of taxa and seminal quality and function parameters. Significant relationships are indicated using p-values. q-values represent Benjamini-Hochberg false discovery rate corrected p-values for multiple comparisons.

| **Sperm quality and function parameters** | **Genera** | **Welch's *t*-statistic** | ***p*-value** | ***q*-value** |
| --- | --- | --- | --- | --- |
| **Sperm DNA fragmentation** | *Finegoldia* | -2.36 | 0.01* | 0.27 |
|  | *Cutibacterium* | -2.20 | 0.02* | 0.27 |
|  | *Porphyromonas* | 2.16 | 0.03* | 0.27 |
|  | *Varibaculum* | 2.11 | 0.03* | 0.27 |
| **ROS** | *Lactobacillus* | 2.18 | 0.02* | 0.66 |
|  | *Corynebacterium* | -2.04 | 0.04* | 0.66 |
| **Semen quality** | *Flavobacterium* | 3.39 | 0.0008*** | 0.02* |
|  | *Prevotella* | 2.26 | 0.02* | 0.38 |
| **Sperm concentration** | *Porphyromonas* | -2.08 | 0.03* | 0.61 |
| **Sperm morphology** | *Flavobacterium* | 3.64 | 0.0003*** | 0.01* |
|  | *Prevotella* | 2.03 | 0.04* | 0.67 |
| **Semen volume** | *Corynebacterium* | 2.27 | 0.02* | 0.32 |
|  | *Actinotignum* | -2.20 | 0.02* | 0.32 |
|  | *Varibaculum* | -2.16 | 0.03* | 0.32 |

**Table III:**
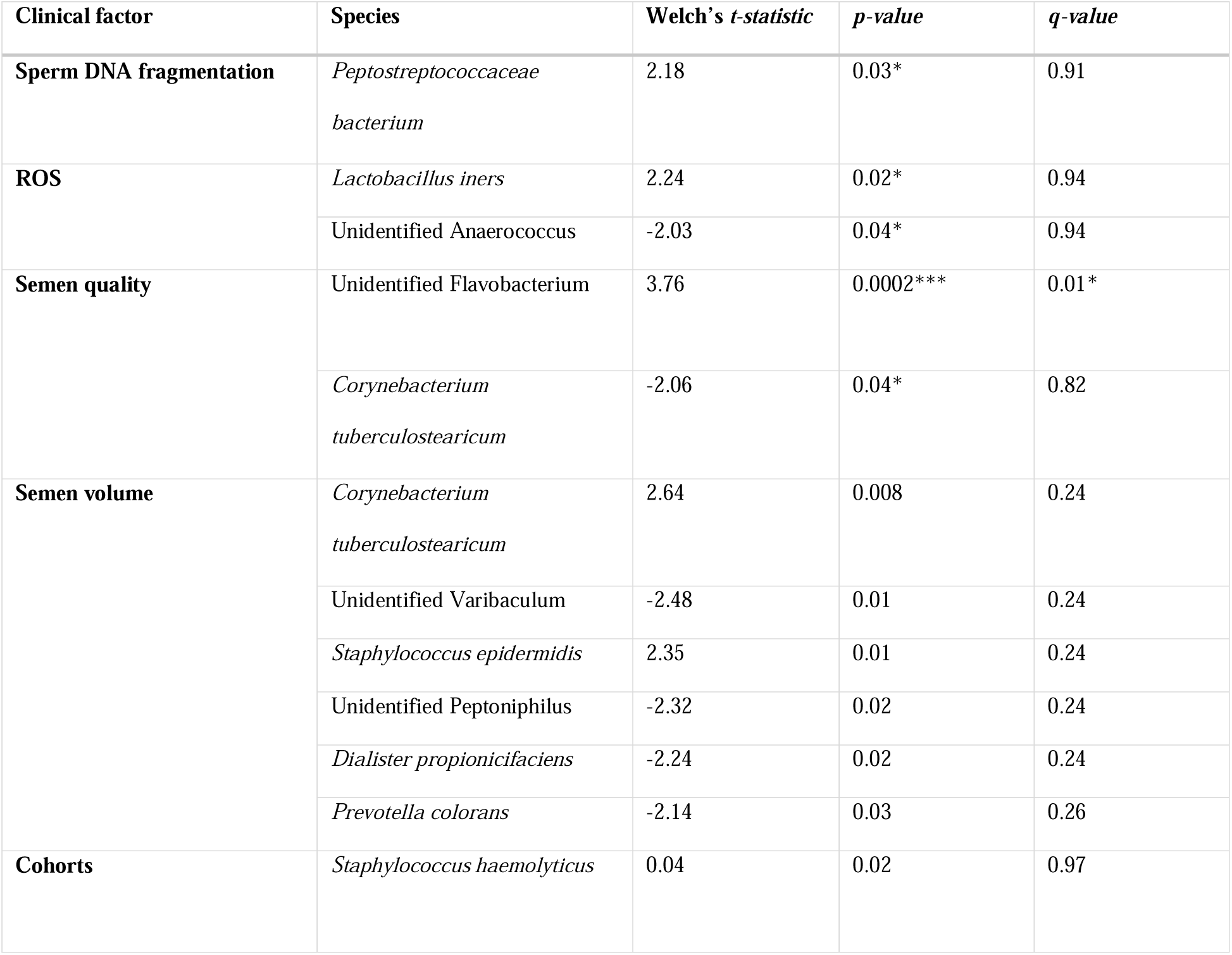
Differential abundance analysis for bacterial species with seminal quality and functional parameters. Positive t-values indicate a positive relationship and a negative t-value describes a negative relationship between relative abundance of taxa and seminal quality and function parameters. Significant relationships are indicated using p-values. q-values represent Benjamini-Hochberg false discovery rate corrected p-values for multiple comparisons.

To focus analyses toward the most extreme phenotype of poor semen quality, a sub-analysis of controls compared with MFI was performed (Table 4). Non-parametric differential abundance analysis again identified a robust relationship between *Flavobacterium* and abnormal sperm morphology (q=0.01, Table 4). At species level, this was mapped to an unidentified species of *Flavobacterium* (q=0.01, Table 5). Similar to findings observed for all samples, sperm DNA fragmentation was inversely associated with relative abundance of *Cutibacterium* and positively associated with *Porphyromonas* and *Varibaculum* was also observed.

**Table IV:** Differential abundance analysis for specific taxa at <u>genera</u> level for controls and cases with male factor infertility. Positive t-values indicate a relationship, and a negative t-value describes a negative relationship between relative abundance of taxa and seminal quality and function parameters. Significant relationships are indicated using p-values. q-values represent Benjamini-Hochberg false discovery rate corrected p-values for multiple comparisons.

| Clinical factor | Genera | Welch's *t*-statistic | *p*-value | *q*-value |
| --- | --- | --- | --- | --- |
| **Sperm DNA fragmentation** | *Cutibacterium* | -2.56 | 0.01* | 0.31 |
|  | *Porphyromonas* | 2.34 | 0.02* | 0.31 |
|  | *Varibaculum* | 1.96 | 0.051 | 0.53 |
| **ROS** | *Finegoldia* | -1.99 | 0.04* | 0.77 |
| **Sperm concentration** | *Finegoldia* | 2.04 | 0.04* | 0.71 |
| **Sperm morphology** | *Flavobacterium* | 3.64 | 0.0003*** | 0.01* |
|  | *Prevotella* | 2.03 | 0.04* | 0.67 |
| **Semen volume** | *Facklamia* | 2.99 | 0.003** | 0.10 |
|  | *Actinotignum* | -2.20 | 0.02* | 0.36 |
|  | *Dialister* | -1.99 | 0.04* | 0.36 |

**Table V:** Differential abundance analysis for specific taxa at species for controls and male factor infertility. Positive t-values indicate a positive relationship and a negative t-value describes a negative relationship between relative abundance of taxa and seminal quality and function parameters. Significant relationships are indicated using p-values. q-values represent Benjamini-Hochberg false discovery rate corrected p-values for multiple comparisons.

| Clinical factor | Species | Welch's *t*-statistic | *p*-value | *q*-value |
| --- | --- | --- | --- | --- |
| **Sperm DNA fragmentation** | *Staphylococcus hominis* | -2.32 | 0.02* | 0.68 |
| **ROS** | Unidentified Flavobacterium | 2.42 | 0.01 | 0.54 |
|  | Unidentified Anaerococcus | -2.12 | 0.03 | 0.54 |
|  | *Schaalia radingae* | -2.12 | 0.03* | 0.54 |
|  | *Haemophilus parainfluenza* | 2.02 | 0.04* | 0.54 |
| **Semen quality** | Unidentified Flavobacterium | 2.36 | 0.01* | 0.91 |
| **Semen volume** | *Dialister micraerophilus* | -2.66 | 0.008** | 0.41 |
|  | *Corynebacterium tuberculostearicum* | 2.27 | 0.02* | 0.44 |
|  | *Staphylococcus epidermidis* | 2.22 | 0.02* | 0.44 |
|  | *Actinotignum schaalii* | -2.00 | 0.04* | 0.45 |
| **Cohorts** | *Staphylococcus haemolyticus* | 0.04 | 0.01* | 0.68 |

## Discussion

We report the largest study to date investigating the seminal microbiota from patients suffering a range of adverse reproductive outcomes including unexplained infertility, male factor infertility and recurrent pregnancy loss. Moreover, we provide a detailed assessment of the relationship between semen microbial diversity, load and compositional structure with both molecular and classical seminal parameters. We identified 3 main clusters present in all study groups. Whilst overall bacterial composition was not association with aberrations in semen analysis, ROS and DNA fragmentation, men with unidentified *Flavobacterium* species were more likely to have abnormal semen quality or sperm morphology.

Several recent studies have indicated the existence of a semen microbiota, however these studies have been limited to small samples sizes and have failed to reach consensus on the compositional structure of the microbiota or its biological relevance, particularly in the context of sperm function and quality (13, 14, 16, 17, 18, 19, 34). By analysing the samples of 233 men with various reproductive disorders we offer a robust assessment of association between the microbiota and classical seminal parameters, but also key functional parameters including seminal ROS and sperm DNA damage. We incorporated stringent negative controls to permit removal of sequences likely originating from extraction kits and reagents known to contaminant low biomass samples such as semen (14, 16, 34). Molina et al report that 50%-70% of detected bacterial reads may be environmental contaminants in a sample from extracted testicular spermatozoa (35); with the addition of passage along the urethra it is likely that contamination of ejaculated semen would be much higher.

Mapping of genera level relative abundance data enabled semen samples to be categorised into 3 major clusters characterised by differing relative abundance of *Streptococcus*, *Prevotella*, *Lactobacillus* and *Gardnerella*. Unlike previous studies, we used an objective statistical approach (i.e. Silhouette methods) to determine the optimal number of microbial clusters supported by the data. These findings are largely consistent with earlier semen metataxonomic profiling studies reporting clusters enriched for *Streptococcus*, *Lactobacillus* and *Prevotella* (13, 14, 16). Moreover, Baud *et al.,* reported increased bacterial richness in the *Prevotella*-enriched cluster, which we also observed (16). This may suggest that certain compositional characteristics of seminal microbiota are conserved across populations. However, similar modelling of species level data, failed to identify statistically robust clusters. This contrasts with other niches such as the vagina where reproducible clusters based on species level metataxonomic profiles have been demonstrated reflecting mutualistic relationships between specific species and the host, which have coevolved over long periods of time (36, 37). It is possible therefore that our findings indicate that microbiota detected in semen are likely the result of transient colonisation events. Consistent with this, several species known to be commensal to the penile skin including *Streptococcus*, *Corynebacterium* and *Staphylococcus*, or the female genital tract including *Gardnerella* and *Lactobacillus*, were observed in semen samples (38). This is in keeping with data suggesting microbiota transference during sexual intercourse (39). It remains possible that a proportion of bacteria detected in semen reflects contamination of the sample acquired during the collection procedure. Studies undertaking assessment of female partner microbiota profiles as well as temporal profiling of semen microbiota would improve understanding of potential dynamic restructuring of semen microbiota compositions. This has been done in part by Baud et al by studying the subfertile couple as a unit to establish if there is a ‘couple microbiota’(40). They took samples from 65 couples with a range of pathologies including idiopathic infertility. From each woman they took vaginal swabs and follicular fluid samples. From each man they took a semen samples and penile swabs. They undertook extensive negative control series and stringent in silico elimination of possible contaminants. The found the male microbiota to be much more diverse than the female, with 90% of female samples being *Lactobacillus*-dominant. Intra-personal male samples i.e. semen and penile swabs from the same man bore more similarity to each other than inter-personal samples of the same sample type i.e. semen *or* penile swab comparisons between men (40). They identified that the male microbiota had very little impact of the microbiota of the female sexual partner(40). Lack of information regarding the sexual activity of the enrolled couples limits this study somewhat.

Several previous studies have described semen microbiota composition to genera level and some have reported associations between specific genera and parameters of semen quality and function (13, 14, 16, 17, 18, 19, 34). However, in many cases these studies have failed to consider multiple comparisons testing, likely leading the reporting of spurious associations. We did not observe any significant associations between bacterial clusters, richness, diversity or load with traditional seminal parameters, sperm DNA fragmentation or semen ROS. This is in contrast with Veneruso *et al.,* who reported that in infertile patients, semen bacterial diversity and richness was decreased whereas Lundy *et al.,* reported that diversity was increased in infertile patients (17, 34). Further, Lundy et al., reported *Prevotella* abundance to be inversely associated with sperm concentration; this was not replicated in our study (17). There are several possible reasons accounting for the high heterogeneity in results including differences in methodology used to assess the microbial component of semen as well as differences in study design (41). For example, time of sexual abstinence prior to sample production as well as sample processing time often differs between studies, which has been shown to impact microbiological composition of semen (42).

The only association between bacterial taxa and semen parameters to withstand false detection rate testing for multiple comparisons detected in our study was between *Flavobacterium* and abnormal semen quality and sperm morphology (q=0.02). The Flavobacterium genus taxon we identified as significantly associated with abnormal semen quality and sperm morphology was present in 36.28% of the samples, with a mean relative abundance of 1.15% in those samples. This information and the mention of previous findings of Flavibacterium in contamination studies have been added to the discussion. *Flavobacterium* are gram-negative physiologically diverse aerobes, some of which are pathogenic (43). *Flavobacterium* was recently identified as a dominant genus in immature sperm cells retrieved from testicular biopsies of infertile men in a study by Molina et al (35). However, in contrast to these findings, a recent smaller study investigating semen collected from 14 sperm donors and 42 infertile idiopathic patients reported an association between *Flavobacterium* and increased sperm motility but a negative correlation with sperm DNA fragmentation (18). The genus Flavobacterium was defined in 1923 to encompass gram-negative, non-spore-forming rods, of yellow pigment (44). The inclusiveness of this definition resulted in a collective of heterogenous species. By 1984 the genus had been restricted to those that were also non-motile and non-gliding (44). More recently, with an increase in genomic profiling, many species previously considered to be of genus Flavobacterium have been reclassified to genus Chryseobacterium, Cytophaga, and Weeksella (45). Increasing numbers of Flavobacterium species are being discovered such as *gondwanense, Collinsii, branchiarum, branchiicola, salegens* and *scophthalmum* (46) (47) (48). The allocation of Flavobacterium *aquatile* to this genus remains controversial due to its motility (49). Flavobacterium species are widely distributed in the environment including soil, fresh water and saltwater habitats (50) (51). There are many reports of pathogenic infections of Flavobacterium species in fish, however human infections are rare (48). A handful of case reports have described opportunistic infections to include pneumonia, urinary tract infection, peritonitis and meningitis (52) (53) (54) (55). Flavobacterium *lindanitolerans* and Flavobacterium *ceti* have been isolated as causative agents in some (56) (54). Case reports also describe Flavobacterium *odoratum* as a causative agent in urinary tract infection, most often in the immunocompromised or those with indwelling devices (57) (58) (59). However, this was one of many species previously of genus Flavobacterium reclassified, in this case to genus Myroides (60). Notably in our sample participants were asymptomatic of urinary tract infection.

Though not withstanding multiple correction, we did observe several other associations between specific bacterial taxa and semen parameters. For example, samples enriched with *Lactobacillus* had lower incidence of elevated seminal ROS, a relationship which could largely be accounted for by *Lactobacillus iners*, a common member of the cervicovaginal niche (61). Various studies have also found *Lactobacillus* enrichment in semen to associate with normal seminal parameters, especially morphology (14, 16). were *Lactobacillus*-predominant (14). However, an association between samples enriched with *Lactobacillus* and asthenospermia or oligoasthenospermia has also been described (19). We also observed an association between increased sperm DNA fragmentation and samples enriched with *Varibaculum*, which is consistent with previous reports of increased relative abundance of *Varibaculum* in semen infertile (34).

A limitation of this and other similar studies is that it was a single institutional study with limited ethnic diversity and potential geographical changes induced by environment or dietary habit. Gut microflora is known to display geographical variability; we cannot exclude that similar geographical variability exists for the seminal microbiota (62). The universal primers used during NGS may not be universal and may anneal variably to specific bacteria resulting in over-detection, under-detection, or indeed non-detection of some taxa (63) (64). A further limitation of this study, and others, is the lack of reciprocal genital tract microbiota testing of the female partners, or paired seminal and urinary samples from male participants. Additionally, we did not have other covariables such as smoking status with which to include in further analyses.

In summary, our study confirms that compositionally the semen microbiota can be broadly classified into three major groups based upon relative abundance of key bacterial genera. Despite different methodological approaches a number of studies, including our own, indicate a Prevotella-dominated or Lactobaccilus-dominated seminal microbiota, perhaps suggesting a stable microbiota at genera level. Our species level our data, however, failed to show similar clusters, perhaps instead suggesting transient colonisation. Longitudinal studies are required to ascertain the stability of the seminal microbiota. We provide evidence for an association between *Lactobacillus* abundance and normal seminal parameters. However, our results indicate that no specific semen bacterial composition can robustly differentiate between fertile and infertile men, although a small subset of bacteria may be associated with changes in seminal parameters. Our finding that an unidentified species of *Flavobacterium* impairs seminal parameters warrants further exploration and may offer the potential for targeted therapies. Larger, multi-centred studies as well as mechanistic investigations are required to establish causal links between the semen microbiota and male fertility.

## Authors roles

Mowla, Farahani, Tharakan, Jayasena and MacIntyre made substantial contribution to the study design, acquisition of data, analysis and interpretation of data and critical revision of the article for important intellectual content. Davies and Correia made substantial contribution to the analysis and interpretation of data and drafting the article. Lee, Kundu, Khanjani, Sindi and Khalifa made substantial contribution to the acquisition of data and critical revision of the article for important intellectual content. Rai, Regan, Henkel, Minhas Dhillo, Ben Nagi and Bennett made substantial contribution to the study design and critical revision of the article for important intellectual content. All authors approved the final version to be published and are in agreement to be accountable for all aspects of the work in ensuring that questions related to the accuracy or integrity of any part of the work are appropriately investigated and resolved.

## Acknowledgements

We would like to thank the patients and participants for their involvement in the study

## Data Availability Statement

The 16S rRNA metataxonomic dataset and the data analysis scripts are publicly available at the European Nucleotide Archive (Project accession PRJEB57401) and GitHub (repository link https://github.com/Gscorreia89/semen-microbiota-infertility), respectively).

