## Supplementary material for "Characterisation and comparison of semen microbiota and bacterial load in men with infertility, recurrent miscarriage, or proven fertility": Supplimental data

**Supplementary Material for Mowla et al., Characterisation and comparison of semen microbiota and bacterial load in men with infertility, recurrent miscarriage, or proven paternity.**

**Supplementary Methods**

**Study design and patient recruitment** This was a case-control clinical study, the experimental arm included men with Male Factor Infertility (MFI), Unexplained Infertility (UI), and male partners of women with Recurrent Pregnancy Loss (RPL). Ethical guidelines for human research were adhered to, and all participants provided written informed consent. Ethical approval was granted by the West London and GTAC Research Ethics Committee (14/LO/1038) and by the Internal Review Board at CRGH (IRB-0003C07.10.19).

Samples were recruited between January 2019 and March 2020. Participants were recruited from The Centre for Reproductive & Genetic Health (CRGH, 230-232 Great Portland Street, London, W1W 5QS), the recurrent miscarriage clinic at St. Mary’s Hospital (Praed street, London W2 1NY). Participants in the control group were recruited through local posters at two hospital sites St. Mary’s Hospital and Hammersmith Hospital, London (72 Du Cane Road, London W12 0HS). Inclusion criteria for both groups were men aged 18-60 years old and BMI < 35kg/m^2^. To be included in the study group participants must have had at least two consecutive miscarriages, and to be included in the control group, the participants must have a history of conceiving and having a child with their partner. Exclusion criteria were a confirmed female cause for RPL, current symptoms of genitourinary tract infections, alcohol excess, hormone therapy, smoking within the last 6 months, and active treatment for severe systemic disease.

**Sample collection** Semen samples were produced by means of masturbation. Patients were asked to wash and dry their hands thoroughly and capture the entire sample in the sterile pots provided. The pots were labelled with patient name, date of birth and other relevant information. Upon completion, they then placed the collection pot inside a hatch from which the laboratory staff collected it. It was then placed in an incubator at 37°C for a minimum of 20 mins prior to analysis and taking an aliquot for storage. The aliquots were collected in sterile cryovials and stored at -80°C until the day of analysis.

**Semen analysis** Semen analysis was carried out by qualified Andrologists in the Andrology Departments of both Hammersmith Hospital and CRGH according to WHO 2010 guidelines and UK NEQAS accreditation (1). Microscopic and macroscopic semen qualities were assessed within 60 mins of sample production. This included: measurement of semen volume, sperm concentration, total sperm count, progressive motility and total motility count, morphological assessment, anti-sperm antibodies and leucocyte count.

**ROS analysis** ROS levels in semen samples was analysed in the Andrology laboratory using an in-house developed chemiluminescence assay validated by Vessey et al. (2). In this assay, Luminol is oxidised in the presence of ROS, which generates chemiluminescence. ROS is measured indirectly by measuring the chemiluminescence generated using a CE marked luminometer (GloMax°Promega Corporation) (2). In advance of ROS measurement, a 100 mmol/L luminol (5-amino-2,3-dihydro-1,4-phthalazinedione; Sigma-Aldrich) stock solution was prepared in dimethylsulfoxide (DMSO). The stock solution generally remains stable for up to 15 weeks. Due to light sensitivity of the assay, the analysis was done in dark and in an aluminium foil covered polystyrene Falcon tube. All reagents stored at 4°C, were brought to room temperature before analysis. A luminol working solution (5 mmol/L luminol prepared in DMSO), negative controls (400μl PBS with 10μl 5 mmol/L luminol working solution), and positive control samples (395μl PBS, 5μl 30% H_2_O_2_ (VWR), and 10μl 5 mmol/L luminol working solution). On the morning of each analysis, the luminometer was calibrated using the negative and positive controls. All samples were mixed gently immediately before placing the tubes in the luminometer. Sperm concentration of >1 M/mL is required for this assay. This is because ROS measurement could be unreliable in lower than 1 M/mL sperm concentrations (3). A total of 400μl of semen was aliquoted using a positive displacement pipette and placed into a 1.5 mL microfuge tube. Following this, 10μl of the luminol working solution was added and gently mixed before taking readings in the luminometer. This process is time sensitive and was done at exactly 20 mins of sample production (minimum amount of time for semen to reach liquefaction). A total of 10 consecutive readings were recorded at 1 min apart. The ROS value was obtained by subtracting control mean from the test mean and correcting it for sperm concentration. Results are therefore reported as ‘relative light units per second per million sperm’. The upper limit of optimal ROS was internally determined at 3.77 RLU/sec/10^6^ sperm (95% CI).

**Sperm DNA fragmentation assessment** for 183 samples TUNEL (Terminal deoxynucleotidyl transferase biotin-dUTP Nick End Labelling) assay (n=183) was used locally. Elevated sperm DNA fragmentation was defined as >20% via TUNEL assay (4). For the remaining 40 samples, Comet assay (a single gel electrophoresis assay) was used. Samples for the COMET assay were sent to the Examen Lab (Belfast, UK) for analysis. Elevated sperm DNA fragmentation was defined as >27% via COMET assay (5).

**Controls and contamination** a series of negative kit/environmental controls were included to identify potential sources of contaminant and elimination of contaminants. Identification and removal of contaminant amplicon sequence variants (ASV) was done using the decontam package (v1.9.0) in R (6).

**DNA extraction** DNA extraction was performed using the QIAamp DNA mini kit as per manufacturer’s instructions (7). samples were removed from -80˚C storage on the day of the analysis and placed on wet ice to thaw. 200μl of semen was used for the process of DNA extraction. Samples were collected in an Ultraviolet (UV) radiated 2ml Eppendorf tube. 300µl of enzymatic master mix including (Lysozyme 10mg/ml, Mutanolysin 25U/μl, Lysostaphin 4000U/ml) was added. After addition of the enzymatic master mix, the samples were pulse vortexed to mix the pellet with the enzymes and incubated at 37˚C for 1 hour. Following the incubation period, samples were pulse centrifuged to bring down any condensation gathered on the inside the lid or on the sides of the tubes. After completion of the enzymatic lysis, the cells in the solution were further disrupted by bead beating. A total of 100mg of bleached and rinsed 0.1mm diameter zirconia/silica was added to each tube and oscillated at 25 Hz for 1 min using a Tissue Lyser. The samples were then pulse spun and 200µl of the solution was transferred to a sterile 1.5ml tube with care taken not to carry forward any beads into the new tube. A total of 20µl of proteinase K was added to each tube along with 200µl of buffer AL and mixed. The samples were then incubated at 56˚C for 30 mins before adding 200µl of absolute ethanol was added and pulse vortex and pulse centrifugation. The solution was then added to a QIAamp (Qiagen) mini spin column in a 2 ml collection tube without wetting the rim of the column. The caps were then firmly shut, and the tubes were centrifuged at 6,200 x g for 1 min. The collection tubes were then discarded, and the columns were placed in fresh 2ml collection tubes. Next, 500µl of AW1 buffer was added without wetting the column rim and the tubes were centrifuged at 6,200 x g for 1 min. Columns were then placed in a fresh collection tube and 500µl of AW2 buffer was added. Column lids were firmly shut, and the tubes were centrifuged at 17,000 x g for 3 mins. The columns were placed in a fresh collection tube and centrifuged at 17,000 x g for 1 min to eliminate the chance of possible AW2 buffer carryover.

The extraction continued by placing the columns in a sterile 1.5µl tube and 100µl of AE buffer was added. The solutions were then incubated at room temperature for 10 mins and further centrifuged at 17,000 x g for 1 min. 5µl of the eluates were aliquoted for PCR to confirm successful extraction. 20µl of the eluates were separately aliquoted, stored at -20˚C and sent for sequencing to Research and Testing Lab in Texas USA, using Illumina MiSeq platform (Illumina, Inc. San Diego, California).

**Quantitative bacteriology** bacterial load was estimated by determining the total number of 16S rRNA gene copies per sample (8). A known concentration of *E. coli* DNA was used to create a ten-fold standard curve from 300 to 30,000,000 copies of 16S DNA. The *E. coli* standards and the 5μl of samples DNA templates were combined with a platinum PCR supermix UDG containing 50nM Rox, BactQuant forward primer at 100μM (5’ CCT ACG GGA GGC AGC A) and reverse primer at 100μM (: 5’ GGA CTA CCG GGT ATC TAA TC), and probe (5’ 6FAM-CAG CAG CCG CGG TA-MGBNFQ) were loaded in duplicates onto a PCR plate. Following a 2 min centrifugation the plate was transferred to the StepOne© qPCR machine. PCR parameters were ser to hold at 50°C for 2 min, 95°C for 10 secs, followed by 40 cycles of 95°C for 15 secs and 60°C for 1 min before a final holding stage of 4°C until plate retrieval. StepOne© software was used to calculate the 16S DNA copy number of the sample in template in relation of the *E. coli* standard curve. Template free PCR controls were included in each run to exclude contamination of PCR reagents.

**16S rRNA gene sequence processing**

Mixed forward primers 28F-YM GAGTTTGATYMTGGCTCAG, 28F-Borrellia GAGTTTGATCCTGGCTTAG, 28F-Chloroflex GAATTTGATCTTGGTTCAG and 28F-Bifdo GGGTTCGATTCTGGCTCAG at a ratio of 4:1:1:1 with 388R reverse primers were used to amplify the V1-V2 hypervariable regions of 16S rRNA gene amplicons. Sequencing was performed on the Illumina MiSeq platform (Illumina, Inc. San Diego, California). Primer sequences were trimmed using cutadapt (9) and read quality was checked using FastQC (10). ASV counts per sample were calculated and denoised using the Qiime2 pipeline (11) and the DADA2 denoising algorithm (12). ASVs were taxonomically classified to species level using a naive Bayes classifier (13) trained on all sequences from the V1-V2 region of the bacterial 16S rRNA gene present in the SILVA reference database (release 138.1) (14) .Extraction kit and negative reagent controls were used to identify and exclude potential contaminants using the decontam package (v1.9.0) in R (15).

**Statistical analysis**

Hierarchical clustering with Ward-linkage and Jensen-Shannon distance was used to assign samples to putative community state types, with the number of clusters chosen to maximise the mean silhouette score. Hierarchical clustering analyses and associated figures were performed in Python (v3.9.12) and with the ‘*pandas*’ (v1.4.2), ‘*scipy’* (v1.7.3), ‘*matptlotlib’* (v3.5.1), and ‘*seaborn’* (v0.11.2) libraries. Principal coordinate Analyses (PCoA) with Jensen-Shannon distance were implemented in R (v4.2.0) using the packages ’*vegdist*’ (v1.40.0) and ‘*ecodist*’ (v2.0.9), and figures generated with *‘ggplot2’* (v3.3.6). Network co-occurrence analyses were also performed in R, with the ‘*SpiecEasi*’ (v1.1.3) implementation of the SparCC method. Species-level count data were filtered by excluding all taxa with a mean relative abundance lower than 1%. SparCC correlations were bootstrapped (10,000 samples), and pairwise correlations with a bootstrapped *p-value* < 0.05, or SparCC ρ < 0.25 were removed. Network visualization and community detection (using the Louvain method) was performed with the ‘*igraph*’ (v2.0.3), ‘*qgraph*’ (v1.9.5), ‘*ggraph*’ (v2.1.0), ‘*tidygraph*’ (v1.2.3) (16). Linear regression models used to regress microbiome features against semen quality parameters and other clinical and demographic variables were fitted with the base R *lm* function. Only features present (non-zero counts) in at least 25% of the samples were carried forward for regression modelling. Centred-log-ratio (clr) transformed (with the `*propr*` (v4.2.6) package) species or genera count values were modelled using linear models with the following formula: *clr*(Species) ~ covariate. For the analyses with estimated absolute species concentrations, log transformation was used instead of the clr transform: *log*(Species + 1) ~ covariate. The ‘*emmeans’* package (v1.7.4.1) was used to perform contrast coding and obtain respective effect size estimates and Welch’s t*-*test *p-values*. Chi-square tests were calculated with the base R *chisq.test* function with Monte Carlo simulation of *p-values* and 1000 resampling replicates (*simulate.p.value=T*, *B=1000*). The Benjamini-Hochberg false discovery rate (FDR) correction (via the *p.adjust* R base function) was used to control the FDR of each covariate signature independently (e.g., ROS, DNA Fragmentation, or Semen quality), with a q < 0.05, or 5%, cut-off, in both the regression and Chi-squared analyses. The ‘*ggplot2’* (v3.3.6), ‘*ggstatsplots*’ (v0.9.4), and ‘*scales*’ (v1.20) packages were used for figure generation.

**Supplementary Methods References**

1.	World Health O. WHO laboratory manual for the examination and processing of human semen. 5th ed ed. Geneva: World Health Organization; 2010.

<http://www.bioinformatics.babraham.ac.uk/projects/fastqc/>

16. Vincent D. Blondel J-LG, Renaud Lambiotte, Etienne Lefebvre. Fast unfolding of communities in large networks. J Stat Mech. 2008:P10008.

**Supplementary Tables**

**Supplementary Table 1: Comparison of mean age and prevalence of ethnicities in study recruitment cohorts.** Ethnicity representation amongst recruited cohorts were not significantly different (p=0.38, Chi-square test). RPL: Recurrent Pregnancy Loss, MFI: male Factor Infertility, UI: Unexplained Infertility.

| **Study cohort** | **Age (mean±SD)** | **Ethnicity** |
| --- | --- | --- |
| **Control (n=63)** | **40.1 ± 8** | 39/63 (62%) Caucasian |
|  |  | 24/63 (38%) non-Caucasian |
| **RPL (n=46)** | **38.2 ± 5** | 35/46 (76%) Caucasian |
|  |  | 11/46 (24%) non-Caucasian |
| **MFI (n=58)** | **36.3 ± 4.5** | 41/58 (70%) Caucasian |
|  |  | 17/58 (30%) non-Caucasian |
| **UI (n=56)** | **37 ± 4.7** | 41/56 (73%) Caucasian |
|  |  | 15/56 (27%) non-Caucasian |

**Supplementary Table 2: Distribution of clinical factors, microscopic seminal parameters, confounding factors, and recruitment cohorts according to genera clusters.** Chi-square tests.

| **Factors** | **Thresholds** | **Cluster 1** | **Cluster 2** | **Cluster 3** | **p-value** |
| --- | --- | --- | --- | --- | --- |
| **DNA frag index** | Low | 60 (53%) | 39 (34%) | 15 (13%) | 0.47 |
|  | High | 37 (45%) | 35 (43%) | 10 (12%) |  |
| **ROS** | <3.77 RLU/s | 74 (52%) | 56 (39%) | 13 (9%) | 0.81 |
|  | >3.77 RLU/s | 19 (58%) | 11 (33%) | 3 (9%) |  |
| **Semen volume** | optimal | 105 (50%) | 80 (38%) | 23 (12%) | 0.12 |
|  | sub-optimal | 8 (53%) | 3 (20%) | 4 (27%) |  |
| **Cohorts** | Control | 36 (57%) | 22 (35%) | 5 (8%) | 0.76 |
|  | MFI | 26 (45%) | 25 (45%) | 7 (10%) |  |
|  | RPL | 23 (50%) | 17 (37%) | 6 (13%) |  |
|  | UI | 28 (50%) | 19 (34%) | 9 (16%) |  |
| **Age** | a. <34 | 30 (61%) | 14 (29%) | 5 (10%) | 0.58 |
|  | b. 34-41 | 59 (48%) | 49 (40%) | 16 (12%) |  |
|  | c. >41 | 24 (48%) | 20 (40%) | 6 (12%) |  |
| **Ethnicity** | Caucasian | 82 (53%) | 57 (37%) | 17 (10%) | 0.58 |
|  | non-Caucasian | 31 (46%) | 26 (39%) | 10 (15%) |  |
| **Concentration** | 1. >15 M/ml | 93 (51%) | 67 (37%) | 22 (12%) | 0.96 |
|  | 2. <15 M/ml | 20 (49%) | 16 (39%) | 5 (12%) |  |
| **Progressive motility** | 1. >32% | 105 (51%) | 78 (38%) | 24 (11%) | 0.67 |
|  | 2. <32% | 8 (50%) | 5 (31%) | 3 (19%) |  |
| **Morphology** | 1. >4% | 37 (50%) | 24 (32%) | 13 (18%) | 0.19 |
|  | 2. <4% | 72 (50%) | 58 (40%) | 14 (10%) |  |
| **Semen quality** | Optimal | 41 (53%) | 24 (31%) | 13 (16%) | 0.17 |
|  | sub-optimal | 72 (50%) | 59 (17%) | 14 (33%) |  |

**Supplementary Table 3: Richness and diversity of seminal bacterial based on seminal quality and function parameters.** Categorical classifications of seminal parameters were based on the clinically defined thresholds. Mann-Whitney tests for all except age. Kruskal-Wallis test was used for age.

| **Factors** | **Richness p-value** | **Diversity p-value** |
| --- | --- | --- |
| **DNA frag index** | 0.68 | 0.89 |
| **ROS** | 0.25 | 0.23 |
| **Semen volume** | 0.54 | 0.85 |
| **Age** | 0.14 | 0.12 |
| **Ethnicity** | 0.31 | 0.24 |
| **Concentration** | 0.79 | 0.66 |
| **Progressive motility** | 0.38 | 0.54 |
| **Morphology** | 0.82 | 0.97 |
| **Semen quality** | 0.74 | 0.90 |

**Supplementary Figures**

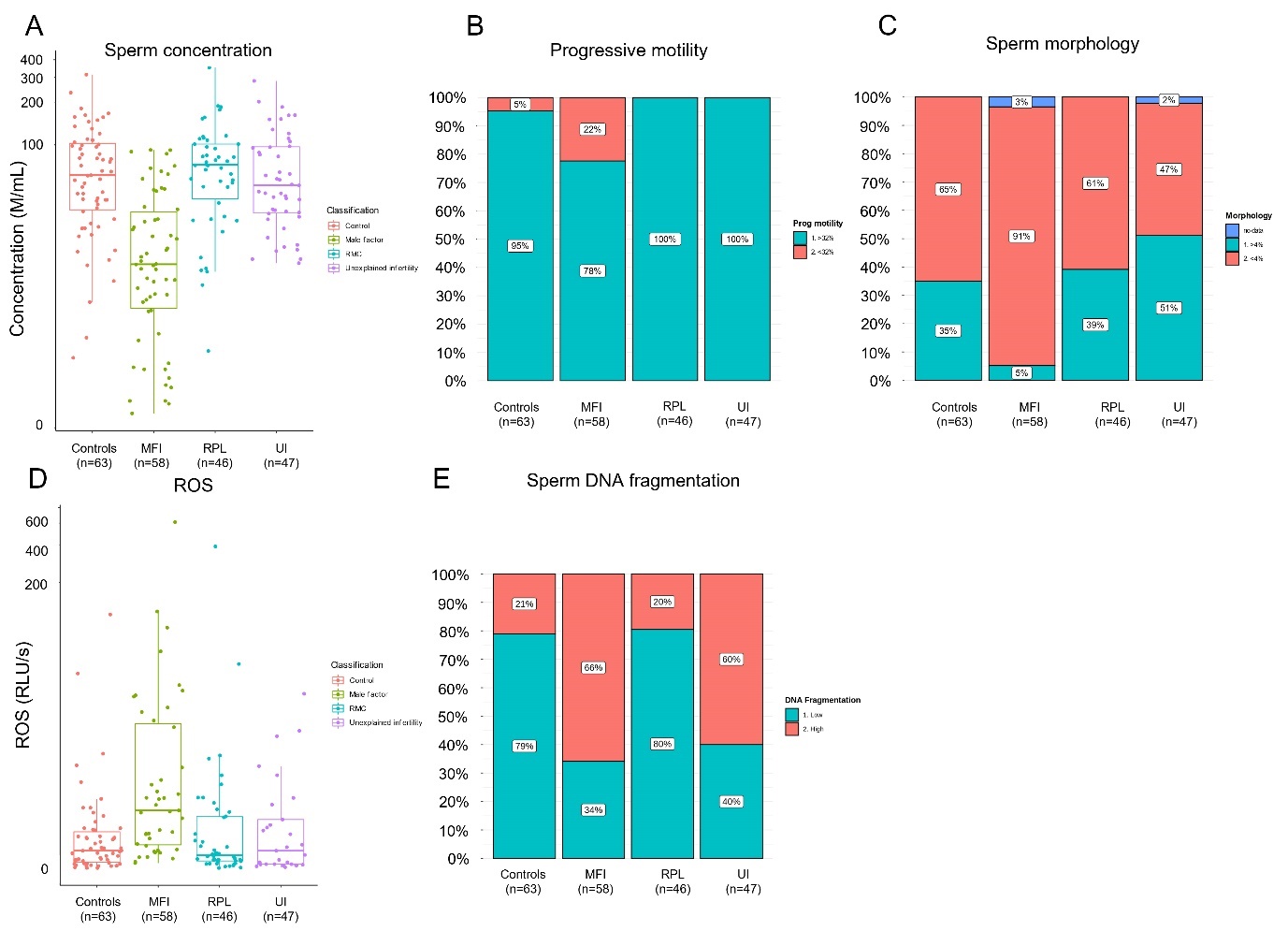

Supplementary Figure 1: Seminal quality and function parameters according to recruited cohorts**.** Comparison of microscopic semen parameters A) concentration (p<0.0001, Kruskall-Wallis rank-sum test) B) progressive motility (p<0.0001, Pearson’s chi-squared test), C) morphology (p<0.0001, Pearson’s chi-squared test) suggested poor semen quality for MFI patients. Comparison of clinical semen qualities D) ROS, E) sperm DNA fragmentation index.

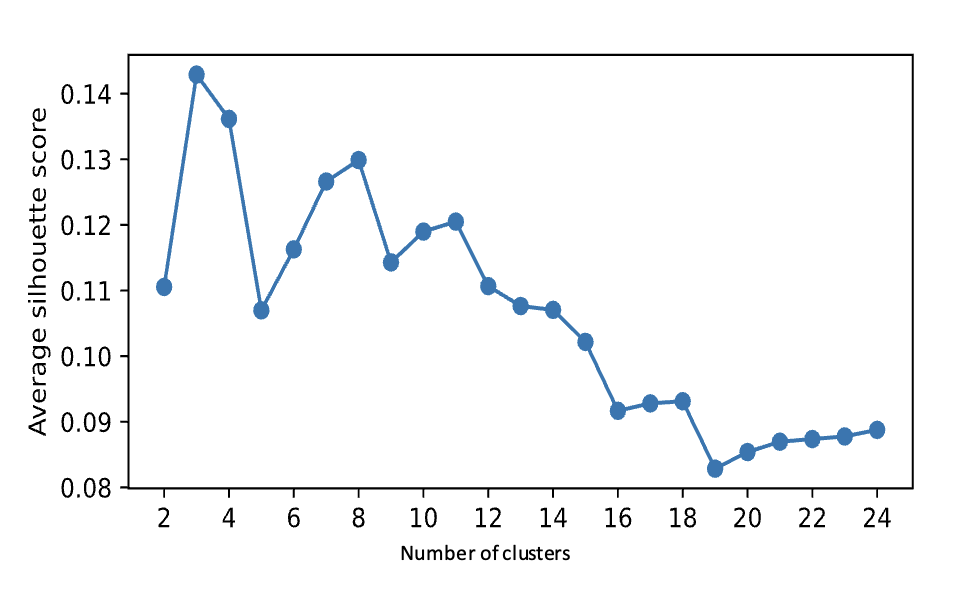

Supplementary Figure 2: Genera level categorization of seminal microbiota identified three major clusters using average Silhouette scores for number of clusters.

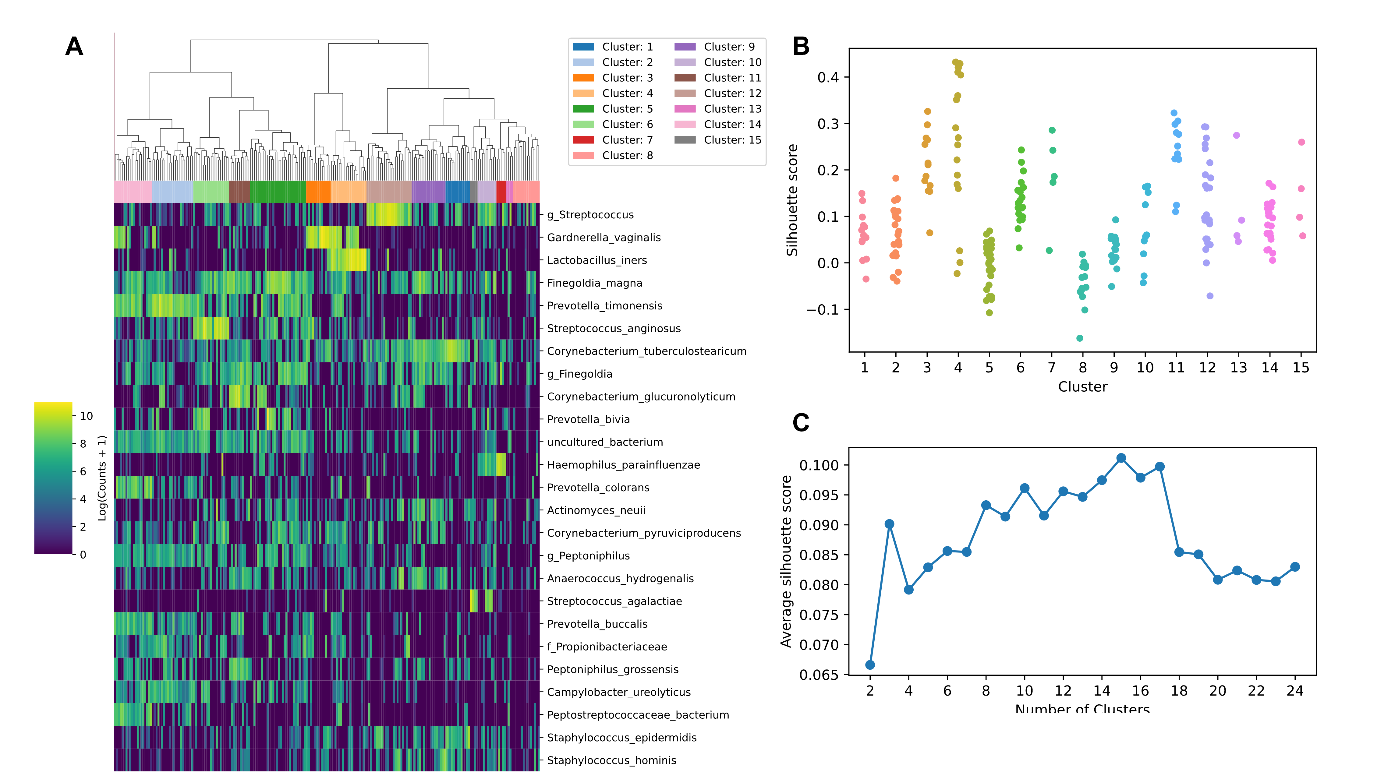

Supplementary Figure 3: Characterization of semen microbiota composition at species level. A) Heatmap of Log10 transformed read counts of top 25 most abundant species identified in semen samples. Samples clustered into 15 microbiota groups. B) Silhouette scores of individual samples in microbial groups. C) Average Silhouette scores for 15 clusters at species level.

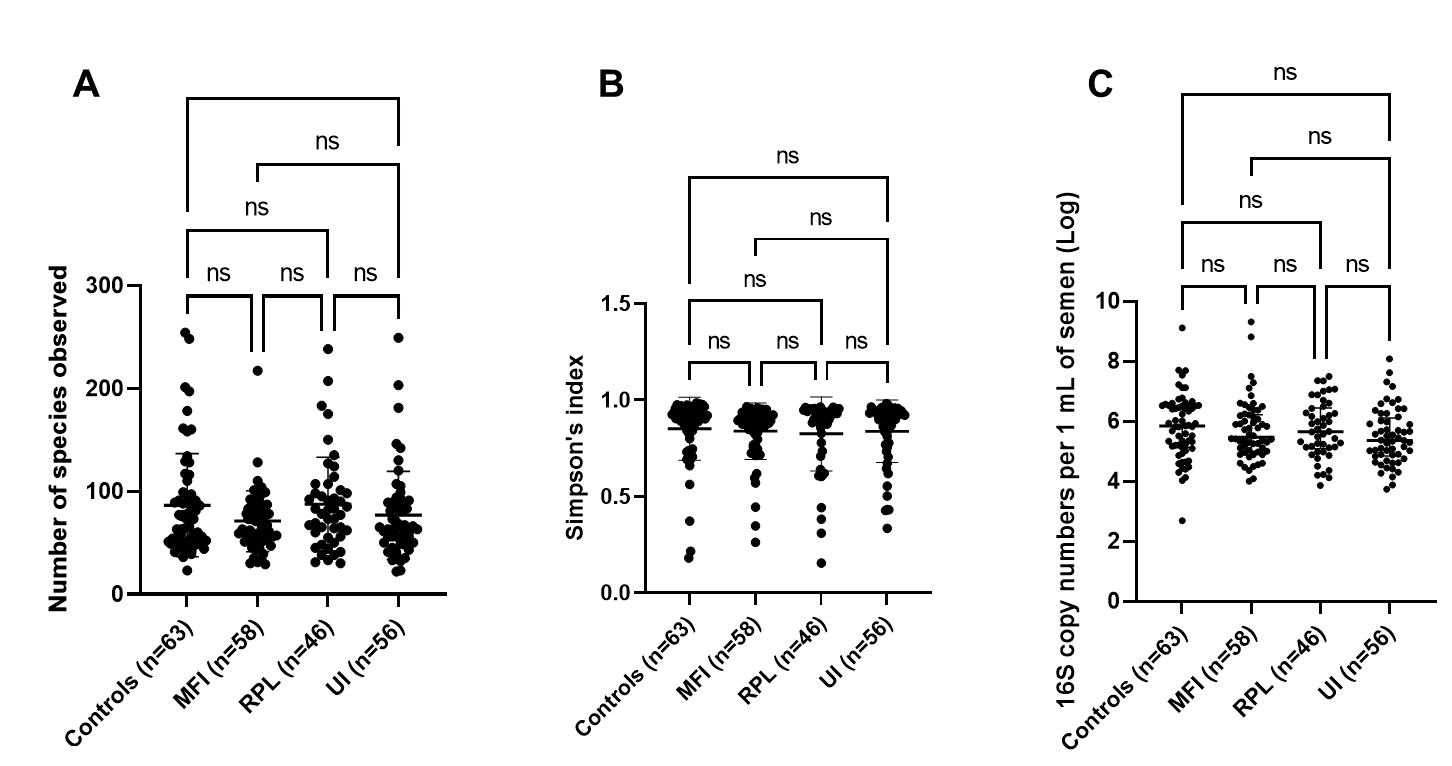

Supplementary Figure 4: **Ecological parameters of seminal microbiota for the recruited study cohorts. A)** Species richness (p=0.30), and **B)** Simpson’s diversity index (p=0.49) were not significantly different based on recruited study cohorts. Kruskal-Wallis tests with Dunn’s multiple comparison p-values demonstrated on the plots. **C)** Bacterial load of seminal microbiota in recruited study cohorts. There were no significant differences in bacterial load based on recruited study cohorts using the number of 16S rRNA genes per 1 mL of semen (p=0.22, Kruskal-Wallis test).
